# Reverse DTNB assay: a novel *in vitro*biochemical approach to detect oxidized thiol modifications

**DOI:** 10.64898/2026.08.02.742231

**Authors:** Ankita Choudhuri, Surupa Chakraborty, Akansha Mishra, Rajib Sengupta

## Abstract

The participation of sulfhydryl or thiol functions in a multitude of protein posttranslational modifications, although reflects on the redox versatility of cysteine residues, but their assessment in a dynamic cellular milieu involving the facile inter-conversion of SH to SSG, S-S, SNO, and S-R’ has been overwhelmingly difficult despite theirimplications in protein folding, enzyme structure and function, signalling and detoxification pathways, and pathophysiological ramifications.The current methodology, in contrast to a wide variety of cumbersome and prolonged techniques,repurposes the conventional DTNB assay for a hassle-free qualitative and quantitative analysisof redox-modified single or multiple susceptible thiol residues of cysteines in pure proteins as well as in a complex mixture of proteins.In this study, we document the thiol content, bearing the susceptibility to undergo reversible, oxidative thiol modifications, utilizing reverse DTNB assay in cell-free lysates and purified proteins that might provide a possible framework for dissecting the physiological phenomena behind the concealment of the susceptible cysteines through their redox-modified forms.

## Introduction

The establishment of redox-based, reversible post-translational modification (PTM) is a direct result of the rapid advent of techniques used in qualitative and/or quantitative estimation of the modified protein thiols (SH) in biological systems. Currently, several methods are capable of selectively recognising oxidative modifications, employing various principles such as direct detection methodologies utilizing radiolabeled(^35^S-)glutathione (GSH), immunoblotting with anti-GSH antibody, tagging or labeling approaches that enrich for S-glutathionylated proteins, or the direct imaging of protein-SSGs within live cells [1,2]. These detection methodologies, in tandem with mass spectrometry that is able to identify the cysteine(s) undergoing S-glutathionylation, intra- or intermolecular disulfide linkages, or S-nitrosylation, have facilitated understanding of the impact this post-translational modification bears on our homeostatic machinery. Additionally, the possibility of oxidative protein post-translational modifications *via* the multiple thiol oxidation pathways that exist owing to the S-glutathionylating capabilities of GSH and its oxidized forms (GSNO and GSSG) makes it imperative to comprehend the existence of differential propensities of the thiol groups of cysteine residues to undergo oxidized thiol modification [3,4]. Thus, in accordance with deciphering the susceptibility of proteins to undergo S-glutathionylation, we employ a novel biochemical approach that utilizes the conventional thiol-quantifying 5,5’-dithiobis(2-nitrobenzoic acid) (DTNB)assay for the quantification of S-glutathionylated thiols, utilizing purified protein as well as a complex mixture of proteins. With the methodology exploiting the thiol-disulfide exchange reaction for the S-glutathionylation of the susceptible thiol and the concomitant release of TNB^2-^(2-nitro-5-thiobenzoate anion),which subsequently aids in the quantification[Figure 1], our results here are indicative of a sensitive qualitative and quantitative determination of redox-modified thiols, demonstrating the susceptibility of proteinthiols towardredox modifications.

**Figure 1:**
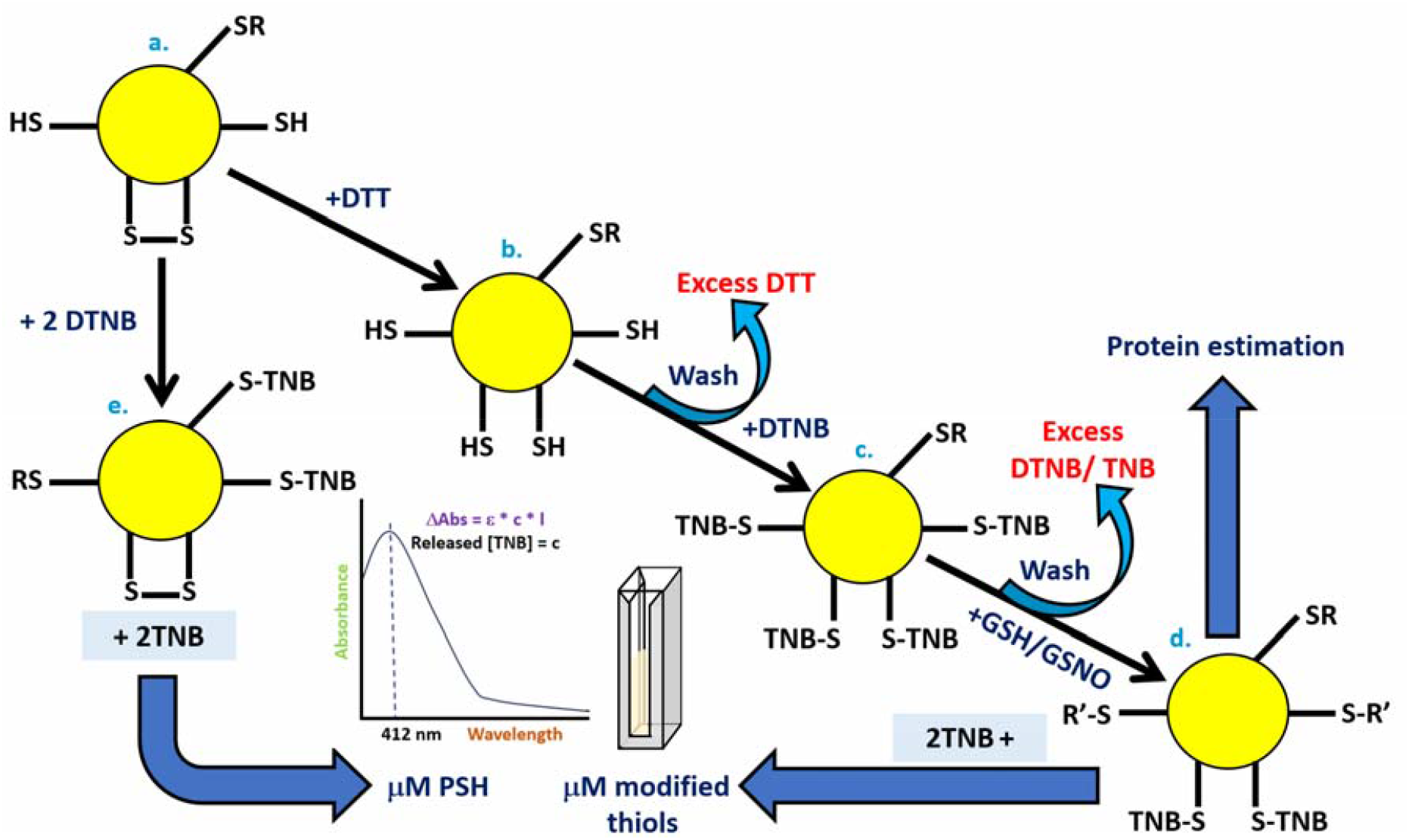
Biochemical analysis of redox-modified protein thiols. S-glutathionylation of susceptible thiols in BSA was observed using a repurposed version of the DTNB assay. Following the reduction of thiol groups by DTT, several washing cycles with a 30 kDa molecular cut-off filter (MCOF) were used to remove any excess, unreacted DTT. This was followed by the addition of DTNB, and subsequent washing steps to remove excess, unreacted DTNB, or TNB^2-^ released from the conjugation of DTNB to the reduced thiols. The attachment of TNB^2-^ to the susceptible thiols, forming a disulfide bond, was exploited in the subsequent step, wherein the addition of reduced GSH to the TNB-bound protein resulted in a thiol-disulfide exchange reaction with the susceptible thiols, leading to the release of TNB^2-^ for every thiol that was added to GSH or GSNO and got modified at single or multiple thiol sites. Spectrophotometric analysis of the released TNB^2-^ at 412 nm provided the quantity of modified thiols (S-thiolation or S-glutathionylation, i.e., RS-SR’, RS-SG, or RSSH formation with GSH or RSNO formation upon GSNO/ S-TNB exchange-mediated S-nitrosylation), and a correlation with the amount of protein present revealed the number of thiols bearing the susceptibility to undergo S-glutathionylation by GSH *in vitro*. (ΔAbs is the absorbance at 412 nm, c is the original concentration, ε is the molar extinction coefficient, 13,600 M^-1^ cm^-1^, l is the path length of the cuvette).

The thiolate/disulfide (RS^−^/RS-SR’, RS^−^/RS-SG, or RS^−^/RSSH) and thiolate/S-nitrosothiol (RS^−^/RSNO) interchanges, briefly elucidated as bimolecular nucleophilic substitution (*S*_N_2) and addition-elimination reaction mechanisms,are of paramount importance in redox biochemistry and can further lead to spatiotemporal and regioselective protein and low-molecular-weight non-protein thiol modifications [Figure 2; 5-8].

**Figure 2:**
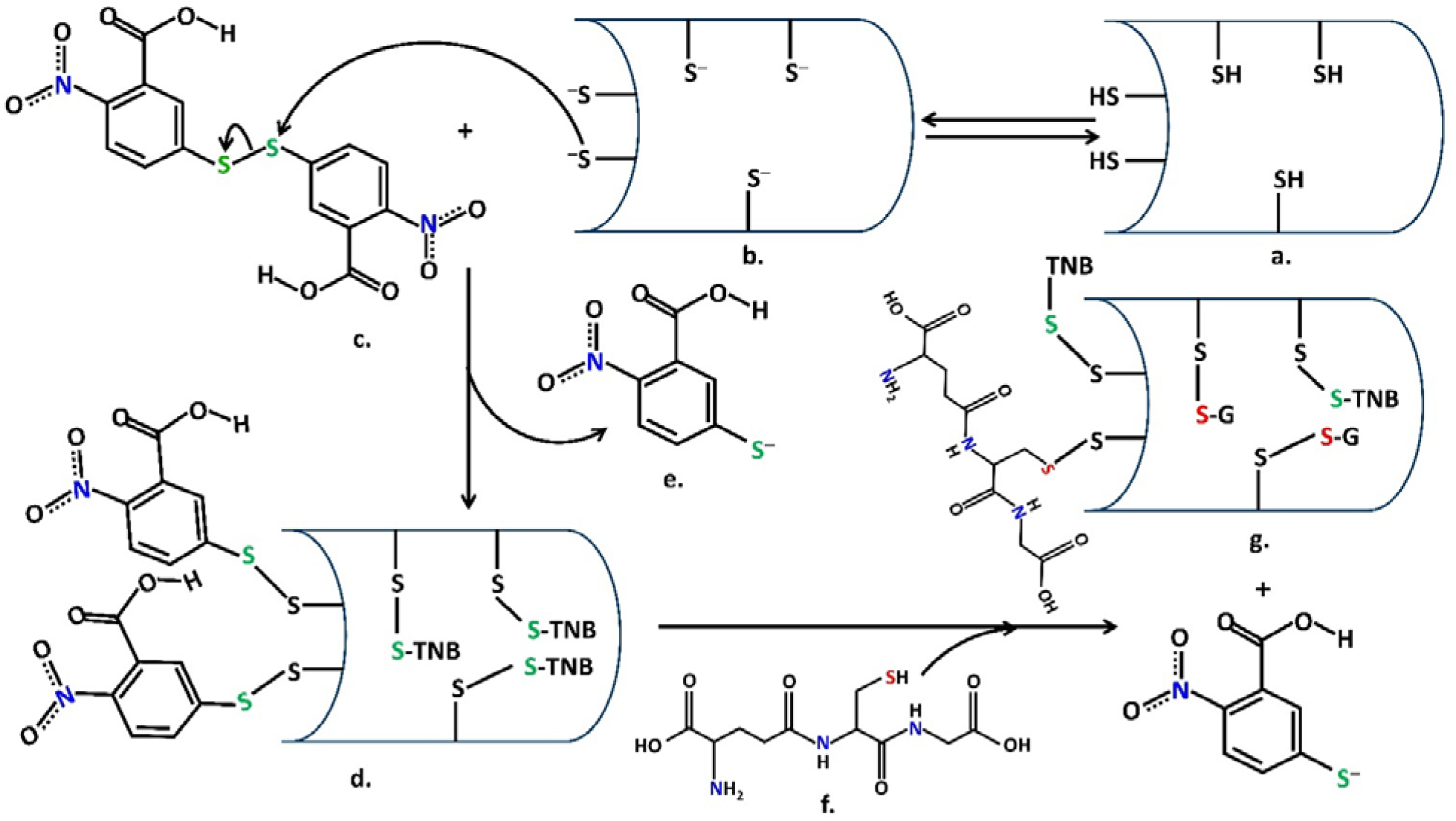
Sequential mechanism of *S*_N_2 thiol-disulfide exchange reaction(s) in reverse DTNB assay. Following DTT-catalyzed pre-reduction of oxidized thiols,a candidate protein harboring protonated (a) or deprotonated(b) reduced free thiols induces thiolate (at low p*K*_a_)-driven nucleophilic attack on the electrophilic disulfide linkage of DTNB (c).TNB-conjugated intermolecular mixed disulfide protein adduct (d) is readily formed as a result of the thiol-disulfide exchange reaction with a concomitant release of the residual TNBanion(one TNB^2-^ released per DTNB molecule), in its thiolate form(e) as a leaving group. Treatment of (d) with physiological levels of low-molecular-weight, reduced glutathione (f) follows another thiol-disulfide exchange reaction, replacing bound TNB anion(s) from thiols susceptible toS-glutathionylation (g).

## Materials and Methods

### Reagents

All experiments were carried out inChelex 100-treated, 0.1 M potassium phosphate buffer (pH 7.4), containing 1 mM EDTA. GSH and BSA were purchased from HiMedia. DTT and DTNB were purchased from Sisco Research Laboratories Pvt. Ltd.

### Reduction of thiols in the sample using DTT

The concentration of DTT used for the reduction of thiols corresponded to the estimated concentration of cysteine residues in BSA. Excess DTT was removed using a 10 kDa molecular weight cut-off filter at 3000 rpm, 4^°^C for 30 minutes. Several washing cycles using the Potassium phosphate buffer described above were carried out till the DTT concentration in the lower fractions was below the detection limit.

### Conjugation of DTNB

DTT-treated samples (after complete removal of DTT) were further treated with DTNB. The concentration of DTNB used for the same was roughly equal to the concentration of thiols in the sample. Excess DTNB was removed using a 10 kDa molecular weight cut-off filter at 3000 rpm, 4ºC, for 30 minutes. The washing cycles with potassium phosphate buffer were carried out till the unreacted DTNB or TNB^2-^ concentration in the lower fractionate was reduced below the detection limit.

### S-glutathionylation and quantification

TNB^2-^-conjugated samples were reacted with 5-10 mM GSH, following which absorbance at 412 nm was taken (ε=13600 M ^-1^ cm ^-1^) to quantify TNB^2-^ released. Protein concentration was determined using the Bradford assay. The concentration of S-glutathionylated thiols in the sample was estimated from the amount of TNB2-released per unit of protein.

## Results and Discussion

The variable number of susceptible thiol residues to either one or more than a single oxidative modification, the high reactivity of -SH, -S-SG, -S-R’, and -S-NO functions, and the short half-life of such reversibly oxidized thiol products in the biological milieu evidently necessitate a rapid, sensitive, reproducible assay readily adaptable to diverse samples for their detection. Research since 1959demonstrated that thiols react rapidly with the chromogenic disulfide, DTNB(Ellman’s reagent), concomitantly releasing the colored anion (product), 2-nitro-5-thiobenzoate anion (TNB^2-^), which corresponds to the SH function of cysteines [9; Figure 1,reaction **a**→**e**]. Utilizing the advantage of DTNB, we examined the quantity of thiols bearing the ability to undergo oxidative modifications in our protein samples (purified proteins and cell lysates), following a pre-reduction step (**a**→**e**), conjugation of reduced free thiols with DTNB to form S-TNB (**b**→**c**), and a thiol-disulfide exchange reaction of **c** with GSH or GSNO (**c**→**d**) [Figure 2]. The simultaneous conjugation of DTNB with pre-reduced thiols and the thiol-disulfide exchange reaction of S-TNB with GSH or GSNO were observed as a means of instantaneous intense yellow coloration in the upper fractionate (qualitative measure) [Figure 1]. The subsequent release of TNB^2-^, as a result of the thiol-disulfide exchange reaction taking place between the TNB-adducted susceptible thiols and GSH/GSNO, was detected by documenting the absorbance at 412 nm using a Hitachi U2910 UV-Vis spectrophotometer, following which protein quantification of the final reaction product was performed using the Bradford assay. Thus, considering the amount of TNB^2-^ released per unit of protein, we determined that, for BSA, approximately ~6 thiols and for mutant C106S DJ-1, about 0.76-1.75 thiols bear the susceptibility to undergo S-glutathionylation with GSH at the endpoint, while in NMuMG cell lysates, about 53.94-75.72 µM/mg/ml of thiols and in *Oryza sativa indica* (rice)-derivedcell lysates, about 46.3 µM/mg/ml were modified upon treatment with GSH or GSNO at thereaction endpoint [Table 1 and 2]. The sequential derivatization of modified thiols from SH largely depends on the thiol susceptibility toward any particular modification [10]. This protocol afforded sensitivity in the micromolar range, relying on the instrumental sensitivity to pinpoint the exact number of thiols that are oxidized in purified proteins and the concentration of oxidized thiols in a cell lysate-derived complex mixture of proteins.

**Table 1:** Estimation of -SR (where R denotes thiol-specific modifications) function upon GSH or GSNO treatment at the endpoint of the reaction mechanism (refer to text for further details). Cell-free lysates were used to perform the assay.

| Sample | -SR content after GSH or GSNO treatment |
| --- | --- |
| NMuMG cell lysate | 75.72 $\mu\text{M}/\text{mg}/\text{ml}$ (+GSH) |
| NMuMG cell lysate | 53.94 $\mu\text{M}/\text{mg}/\text{ml}$ (+GSNO) |
| Rice (*O. sativa*) cell lysate | 46.3 $\mu\text{M}/\text{mg}/\text{ml}$ (+GSH) |

**Table 2:** Estimation of -SR (where R denotes thiol-specific modifications) function upon GSH or GSNO treatment at the reaction endpoint (refer to text for further details). Recombinant purified proteinswere used to perform the assay.

| Sample | Thiol content | Number of cysteines in free thiol form | -SR content after GSH or GSNO treatment |
| --- | --- | --- | --- |
| BSA | 35 | 1 | 6.02 (+GSH) |
| DJ-1 C106S | 2 | 2 | 1.75 (+GSH) |
| DJ-1 C106S | 2 | 2 | 0.76 (+GSNO) |

The concept of S-glutathionylation of proteins encompasses a large spectrum of research across various categories, such as deciphering cysteines undergoing this PTM across different *invitro* and *invivo* contexts; investigating the corresponding changes in the protein structure and function and/or impact on cellular machinery; implementing dry lab techniques to determine the propensity of thiol groups of cysteines within a protein to be S-glutathionylated; and different protocols documenting methodologies that have helped establish this fundamental post-translational modification. One of the most common ways this post-translational modification has been documented across different physiological contexts is by Western blot analysis with anti-GSH antibodies, which can be preceded by immunoprecipitation of the target protein [2,11]. However, the cumbersome nature of Western blot analysis, coupled with the limited sensitivity and specificity of the anti-GSH antibody [1], raises questions regarding the level of sensitivity this methodology is equipped with. Other detection techniques include the use of radiolabeled (^35^S-) GSH by the introduction of ^35^S-cysteine to cell culture for the analysis of *in vivo*SSG; however, it is accompanied by the limitations of false positives generated by radiolabeled cysteinylated proteins and blocking protein synthesis as necessitated by the methodology [1]. However, in recent years, there has been a preference of transitioning to methodologies that rely on tagging or labeling approaches, which are either based on blocking non-modified thiols and then selectively modifying the S-glutathionylated thiol or employing GSH analogs that can directly react with the susceptible thiol [1]. These methodologies significantly overcome the limited sensitivity and specificity of a protocol only relying on the anti-GSH antibody for the detection of S-glutathionylation but are accompanied by the restrictions of drawbacks pertaining to the accessibility and experimental costs. Mullen et al. [12] developed a ‘redox array’ pertaining to the detection of S-glutathionylation, utilizing a unique, combinatorial approach that was based on the direct incorporation or the direct introduction of cell-permeable BioGEE into THP-1 cells, following which the detection methodology eventually employed an antibody-based array for identifying S-glutathionylated proteins. However, the restriction in the number and identity of antibodies, in conjunction with the possible generation of false positives due to the non-specific binding of unmodified proteins, makes it pertinent to cross-check the S-glutathionylated status with other methodologies [12]. The advent of dry lab-based techniques has also ushered in the era of determining the propensity of specific cysteine residues to undergo S-glutathionylation [13]. Thus, accounting for all the advantages and drawbacks, the addition of a novel methodology for detecting S-glutathionylation to this existing repertoire of techniques should only enhance the spectrum of methodologies that can be used for examining this PTM.

Our findings here are limited to the category of *in vitro* determination of protein thiols susceptible to redox modifications, such as S-glutathionylation, utilizing a methodology that repurposes the traditional thiol-quantifying DTNB assay, introducing a novel and economical addition to the existing methodologies. Thus, our results [Table 1 and 2]document the capability of ~6thiols in BSA and 0.76-1.75 thiols in single mutant C106S DJ-1to undergo S-glutathionylation, opening a new dimension forexamining the capabilities of otherwise engaged thiols to undergo post-translational modifications. Conventionally, the S-glutathionylation of BSA has often served as a standard for the assessment of protocols or methodologies targeting the detection of S-glutathionylation of other target proteins, without much insight into the susceptibility of constituent cysteines within BSA to undergo this post-translational modification. Giustarini et al. in their work demonstrated that the prominent S-glutathionylating agents of GSSG and GSNO were unable to S-glutathionylate BSA significantly [14]. In our study, we present evidence of BSA undergoing S-glutathionylation on multiple cysteines by GSH, thus demonstrating the propensity displayed by these cysteines to undergo S-glutathionylation. As documented, out of the 35 cysteines in BSA, Cys34 is said to be in its free thiol form, while the other 34 remain engaged in 17 disulfide bonds. While our methodology does utilize DTT to pre-reduce any reversibly modified thiols and then proceed to the detection methodology, our results, in retrospect, raise questions regarding the susceptibility of these otherwise disulfide-bound cysteines to undergo post-translational modifications, hinting at a possible cellular rationale in keeping these cysteines disulfide-bound in physiological conditions. As assessed by the same assay, a lyophilized, ultrapure C106SDJ-1 mutant, reconstituted in Chelex100-treated phosphate buffer, supplemented with 1 mM EDTA, revealed the propensity of1.75thiol functions with GSH at the reaction endpoint (glutathionylated DJ-1, 49.1 µM; 0.56 mg/mlor 28 µM estimated protein concentration)per mole of protein toward S-glutathionylation. Incubation of TNB-conjugated mutant DJ-1 with GSNO (100 µM) in comparison with an initial pre-reduced (DJ-1-(SH)_2_, 98.2 µM) thiol-enriched DJ-1 fractionresulted in the S-glutathionylation of approximately 0.76 thiol functions(mutant DJ-1-SSG, 5.36 µM; 0.14 mg/ml or 7 µM estimated protein concentration), indicating that GSH, if used at the terminal reaction of this assay, canefficiently catalyze the thiol-disulfide exchange reaction to a significant extent in comparison to GSNO. The preference of GSH or its S-nitroso derivative, GSNO, at the reaction endpoint strongly determines the reaction outcome of modified or oxidized protein thiols, as GSNO, in contrast to GSH, might not necessarily be limited to catalyzing S-glutathionylation but also exertsa potential transnitrosylase activity on susceptible thiols upon -S^−^/-SNO exchange reactions with TNB-bound protein [15-17]. While thiol-disulfide interconversion reinforces the possibility of reversible posttranslational modifications at susceptible cysteine residues [10,18], its principle being repurposed towards designing an *in vitro* reverse DTNB assay system for the detection of modified thiols extends the versatility of biological thiols as reductants *via*non-protein and/or protein thiol or thiolate interactions.

## Author contributions

R.S. conceived the project idea, supervised the project, conducted the experiments, and performed data analyses. S.C., A.C., and A.M.conducted the experiments and data analyses.R.S., S.C., and A.C. wrote the manuscript and revised it.

## Acknowledgements

This work is dedicated to the memories of Dr.Detcho A. Stoyanovsky and Dr. Arne Holmgren. We would like to thank them for their enormous help and support. We are grateful to Dr.Stoyanovsky and Dr. Holmgren for showing us the true meaning of science in their own unique ways. This research was partially supported by Karolinska Institute (Forskningsstiftelser research grant no 2013fobi37677).The authors of the manuscript are grateful to Dr. SantanuPalchaudhuri (Amity University, Kolkata) for kindly providing us with the NMuMG cell line and Professor Mark Wilson (Department of Biochemistry and the Redox Biology Center, University of Nebraska) for the ultrapure C106S DJ-1 protein.We would like to thank Dr.Surajit Bhattacharya (Amity University, Kolkata) for providing the crude cell-free root extract of *Oryza sativa*.

## References

1. Li X, Zhang T, Day NJ, Feng S, Gaffrey MJ, Qian WJ. Defining the S-Glutathionylation Proteome by Biochemical and Mass Spectrometric Approaches. Antioxidants (Basel). 2022 Nov 17;11(11):2272. doi: 10.3390/antiox11112272.

2. Butturini E, Boriero D, Carcereri de Prati A, Mariotto S. Immunoprecipitation methods to identify S-glutathionylation in target proteins. MethodsX. 2019 Sep 10;6:1992–1998. doi: 10.1016/j.mex.2019.09.001.

3. Lermant A, Murdoch CE. Cysteine Glutathionylation Acts as a Redox Switch in Endothelial Cells. Antioxidants (Basel). 2019 Aug 16;8(8):315. doi: 10.3390/antiox8080315.

4. Musaogullari A, Chai YC. Redox Regulation by Protein S-Glutathionylation: From Molecular Mechanisms to Implications in Health and Disease. Int J Mol Sci. 2020 Oct 30;21(21):8113. doi: 10.3390/ijms21218113.

5. Singh R, Whitesides GM. Degenerate intermolecular thiolate-disulfide interchange involving cyclic five-membered disulfides is faster by .apprx.103 than that involving six-or seven-membered disulfides. J Am Chem Soc. 1990 Aug;112(17):6304–9. doi:10.1021/ja00173a018.

6. Bachrach SM, Woody JT, Mulhearn DC. Effect of ring strain on the thiolate-disulfide exchange. A computational study. J Org Chem. 2002 Dec 13;67(25):8983–90. doi: 10.1021/jo026223k.

7. Bach RD, Dmitrenko O, Thorpe C. Mechanism of thiolate-disulfide interchange reactions in biochemistry. J Org Chem. 2008 Jan 4;73(1):12–21. doi: 10.1021/jo702051f.

8. Barnett DJ, McAninly J, Williams DLH. Transnitrosation between nitrosothiols and thiols. J Chem Soc, Perkin Trans 2. 1994;(6):1131. doi:10.1039/p29940001131.

9. Ellman GL. Tissue sulfhydryl groups. Arch BiochemBiophys. 1959 May;82(1):70–7. doi: 10.1016/0003-9861(59)90090-6.

10. Chakraborty S, Choudhuri A, Mishra A, Sengupta R. S-nitrosylation and S-glutathionylation: Lying at the forefront of redox dichotomy or a visible synergism? BiochemBiophys Res Commun. 2025 May 1;761:151734. doi: 10.1016/j.bbrc.2025.151734.

11. Medvedeva MV, Kleimenov SY, Samygina VR, Muronetz VI, Schmalhausen EV. S-nitrosylation and S-glutathionylation of GAPDH: Similarities, differences, and relationships. BiochimBiophys Acta Gen Subj. 2023 Sep;1867(9):130418. doi: 10.1016/j.bbagen.2023.130418.

12. Mullen L, Seavill M, Hammouz R, Bottazzi B, Chan P, Vaudry D, Ghezzi P. Development of ‘Redox Arrays’ for identifying novel glutathionylated proteins in the secretome. Sci Rep. 2015 Sep 29;5:14630. doi: 10.1038/srep14630.

13. Anashkina AA, Poluektov YM, Dmitriev VA, Kuznetsov EN, Mitkevich VA, Makarov AA, Petrushanko IY. A novel approach for predicting protein S-glutathionylation. BMC Bioinformatics. 2020 Sep 14;21(Suppl 11):282. doi: 10.1186/s12859-020-03571-w.

14. Giustarini D, Milzani A, Aldini G, Carini M, Rossi R, Dalle-Donne I. S-nitrosation versus S-glutathionylation of protein sulfhydryl groups by S-nitrosoglutathione. Antioxid Redox Signal. 2005 Jul-Aug;7(7-8):930–9. doi: 10.1089/ars.2005.7.930.

15. Singh SP, Wishnok JS, Keshive M, Deen WM, Tannenbaum SR. The chemistry of the S-nitrosoglutathione/glutathione system. Proc Natl Acad Sci U S A. 1996 Dec 10;93(25):14428–33. doi: 10.1073/pnas.93.25.14428.

16. Broniowska KA, Diers AR, Hogg N. S-nitrosoglutathione. BiochimBiophys Acta. 2013 May;1830(5):3173–81. doi: 10.1016/j.bbagen.2013.02.004.

17. Rossi R, Lusini L, Giannerini F, Giustarini D, Lungarella G, Di Simplicio P. A method to study kinetics of transnitrosation with nitrosoglutathione: reactions with hemoglobin and other thiols. Anal Biochem. 1997 Dec 15;254(2):215–20. doi: 10.1006/abio.1997.2424.

18. Zeida A, Radi R. Thiol oxidation mechanisms and specificity. In: Alvarez B, Comini M, Salinas G, Trujillo M,eds. Redox Chemistry and Biology of Thiols, 1st ed., Academic Press, New York, 2022, pp. 99–113.

